# The conversion of forests to agricultural land reduced the content of soil black carbon fractions in a karst rocky desertification area of Southwest China

**DOI:** 10.1101/2024.09.21.614285

**Authors:** Denan Zhang, Qiumei Teng, Kechao Huang, Yuyi Shen, Yingjie Sun, Shihong Lyu, Guangping Xu, Yanzhao Zhang

**Author notes:** Correspondence (X). These authors contributed equally to this work (Denan Zhang, Qiumei Teng).

## Abstract

Understanding the effects of land use on soil black carbon (BC) is critical for correctly interpreting the role of BC in the carbon cycle of karst areas. This study investigated the distribution characteristics of organic carbon (OC), BC, char, and soot and soil physicochemical properties of the soil profiles (0–40 cm) of secondary forest, shrub, farmland, and wasteland in a rocky desertification area of Guangxi Province, a typical karst rocky desertification area in Southwest China. We used a combination of soot/char and δ^13^C_BC_ isotope analysis to identify the source of BC of these soils. The average value of soil BC was the highest in secondary forest, followed by shrub, farmland, and wasteland in the 0–40 cm soil layer. BC had a positive correlation with OC, char, soot,soil nitrogen, phosphorus and potassium, showed a negative correlation with bulk density. In the black carbon component, char occupies a greater proportion than soot, with the heavy fraction OC exhibiting a higher concentration of BC than the light fraction OC. Conclusively, the BC content was mainly due to C3 plant burning, vehicle exhaust emissions, and fossil fuel utilization.Vegetation restoration improved the soil BC content associated with OC sequestration in the typical karst rocky desertification mountainous area of Southwest Guangxi.

## 1. Introduction

Organic carbon (OC) is a highly complex and heterogeneous mixture of readily decomposable reactive OC and refractory inert OC [1]. Inert OC has a slow decomposition rate, a long turnover time, and relative stability and plays an essential role in the long-term sequestration of soil OC (SOC). Black carbon (BC) is an important component of OC and is generally defined as a solid organic material rich in carbon that results from the incomplete combustion of carbonaceous fuels, such as fossil fuels and biomass. BC consists of a complex mixture of slightly charred biomass compounds and highly condensed refractory materials, including soot, char, charcoal, which are widely distributed in various carriers such as soil and atmosphere. BC has a highly concentrated aromatized structure [2], exhibits high oxidation resistance, is not easily degraded by microorganisms [3], and facilitates soil carbon storage in the long term [4].

As an important component of the inert SOC pool [3], BC plays a crucial role in the global carbon cycle [5]. Global annual BC production is as much as 62–294 Tg, of which 80%–90% is deposited directly into the soil [6]. The use of BC is a critical strategy for climate change mitigation [7] and constitutes a vital component of the “missing carbon” in Earth’s carbon balance [3, 8], because BC can increase carbon sequestration [4], mitigate greenhouse gas emissions [9], and enhance soil fertility [10]. In the literature, BC has been explored in many areas, including forests [11–12], agricultural fields [13], wetlands [14], and urban areas [15]. The results in the literature have demonstrated that BC can effectively increase stable soil carbon stocks and significantly contribute to the mitigation of the greenhouse effect.

Land use has been seriously influenced and disturbed by human activities and is one of the most critical drivers influencing the accumulation or decomposition rate of SOC [16]. Land use also has the potential to modify global carbon cycles and influence climate, as well as influence the accumulation of BC in soils [17]. Rational land use patterns can improve soil structure and resistance. Research on various land use patterns in karst mountains has revealed that SOC is the primary determinant of soil nutrient levels [18], deforestation and cultivation significantly increase soil capacity [19], and land use patterns affect the distribution of OC within the soil profile [20]. Notably, applying an appropriate land use strategy can enhance land use efficiency and contribute to the comprehensive prevention of rock desertification.

A representative area for various land use patterns is in Southwest Guangxi, China. This typical region has widespread karst crest depressions characterized by severe rock desertification and a poor ecological environment [21]. Over the past twenty-two years in Southwest Guangxi, vegetation has gradually been restored in some areas due to the implementation of the Grain-for-Green project [22] and the vigorous implementation of ecological restoration and reconstruction engineering measures in its rocky desertification area. Existing research has focused on soil microorganism quantity and activity characteristics [23], soil physicochemical properties [24], the impact of vegetation on water cycle processes [25], changes in vegetation cover [26], soil enzyme activity characteristics under different land use patterns [27], and carbon sequestration in stone desertification management [28].He et al [29] studied SOC and carbon accumulation in the karst forests of the Guizhou autonomous region. Southwest few studies have focused on the distribution and storage of soil BC, char, and soot in this region.

Land use conversions from forest land to agricultural land (cropland and grassland), which are typical land use conversions that can influence soil carbon stocks [30]. Land use is an important factor controlling SOC fractions, and because of traditional agriculture and different land use patterns, soils may exhibit unique distribution patterns of BC and its sources. Thus, to establish a scientific foundation for evaluating soil quality, this study investigated the impact of human activities on soil BC accumulation during ecological restoration, providing an examination of the effects of different land use patterns on soil BC in Southwest Guangxi’s rocky desertification areas that is of substantial practical importance. We used the rocky desertification area of Southwest Guangxi as our research object, examining soils collected from four land use patterns: secondary forest, shrub, farmland, and wasteland.

The objectives of this study were to 1) investigate the vertical distribution characteristics and potential sources of soil BC and the correlation between soil BC and SOC under varying land use patterns, 2) elucidate the factors correlated between BC and OC in the rocky desertification region of Southwest Guangxi, and 3) evaluate the influence of land use patterns on the spatial distribution and source apportionment of soil BC composition. The overall aim was to provide fundamental data on the soil carbon pool of karst ecosystems in Southwest Guangxi and offer theoretical references for rational soil utilization and environmental protection in rocky desertification areas.

## 2 Materials and Methods

### 2.1 Study area

The study area was located in Guohua Town, Pingguo City, Guangxi Province (107^°^22^′^40^″^–107^°^25^′^30^″^E, 23^°^22^′^30^″^–23^°^24^′^00^″^N), a typical karst area with an elevation of 110 –570 m. The average annual temperature in this area is 19.1–22.0 °C, and the annual precipitation is approximately 1500 mm. The seasonal precipitation distribution is uneven: approximately 70% of the total annual precipitation occurs between May and August, and 30% occurs between September and April.

The soil was predominantly composed of brown lime and was characterized by bare rocks and shallow soil. The vegetation cover was low, and the rocky desertification was serious. The dominant vegetation communities were secondary forest, shrub, and cultivated species. The dominant tree species in secondary forests were *Zenia insignis*, *Melia azedarach*, *Apodytes dimidiata,* and *Choerospondias axillaris*. The main shrub species were *Alchornea trewioides*, *Cipadessa cinerascens,* and *Vitex negundo* [31].

Land use patterns were classified as wasteland, farmland, shrub, and secondary forest, as aforementioned (Fig 1). The cultivated land in the region, typically consisting of maize, soybeans, and pitaya, has been cultivated for at least 110 years. Shrub and secondary forest were estimated to be 39 and 80 years old, respectively, followed by cultivated land abandonment at approximately 17 years.

**Fig 1.**
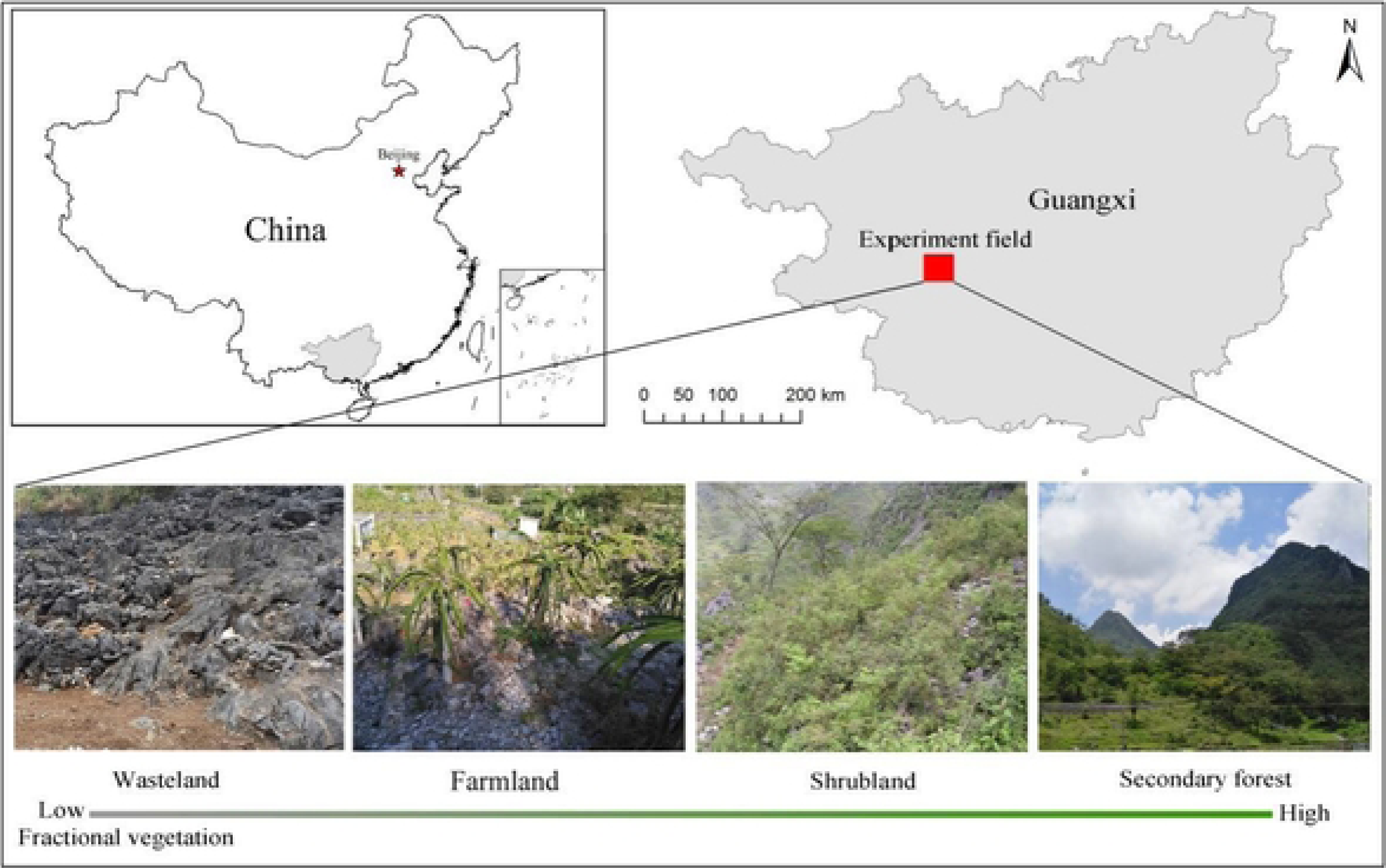
Location map of the study area, with different land use types associated with vegetation restoration.

### 2.2 Soil sampling

Four sampling sites representing different land use patterns (wasteland, cultivated land, shrub, and secondary forest) were carefully selected based on the degree of vegetation restoration. In November 2020, within each sampling site, three replicate plots measuring 50 × 50 m were selected for a total of 12 plots used for soil sample collection. Soil samples were collected from four depths: 0–10 cm, 10–20 cm, 20–30 cm, and 30–40 cm using a soil auger (5 cm in diameter), and soil samples of the same layer were mixed as one soil sample. The presence of anthropogenic management measures, such as fire and fertilization, at each site was investigated. The collected soil samples were placed in sterile self-sealing bags, quickly refrigerated in sealed ice bag containers, and transported to the laboratory for storage at 4 °C in a refrigerator. After the samples were air-dried, all samples were sieved through a mesh of ө < 2 mm and were stored in sealed bottles for analysis.

### 2.3 Sample analysis

#### 2.3.1 BC measurement

This study used a methodology refined from that of Lim & Cachier [32]In brief, the six-step pretreatment proceeded as follows: ① 3 g fine dry soil (ө < 2 mm) was weighted in a glass recipient, ② slowly covered with 15 ml 3 mol·L^-1^ HCl to remove carbonate, and left to react for 24 h; ③ 15 ml 10 mol·L^-1^ HF: 1 mol·L^-1^ HCl was added to remove silicate and left to react for 24 h; ④ it was slowly covered with 15 ml 10 mol·L^-1^ HCl and left to react and remove the possible generation of CaF_2_ for 24 h; ⑤ 15 ml 0.1 mol·L^-1^ K_2_Cr_2_O_7_:2 mol·L^-1^ H_2_SO_4_ was added to remove the OC and left at 55°C±1°C for 65 h; ⑥ the remaining material was the BC sample, which was centrifuged, dried, and analyzed using elemental analysis-isotope mass spectrometry (FLASH EA∼DELTA V) to determine the BC content and the BC stable isotope ratio (δ^13^C PDB).

Referring to the BC method of Zhan et al [33], the temperature was increased in stages to 140, 280, 480, and 580 °C to produce four OC fractions: OC1, OC2, OC3, and OC4. Gas mixture of 2% O_2_/98% He is introduced, the temperature was increased in stages to 580, 740, and 840 °C, producing three elemental carbon fractions: EC1, EC2, and EC3. This study defined this point as the demarcation point between organic and BC and defined the part of EC as pyrolytic carbon (POC). The OC was defined as OC1 + OC2 + OC3 + OC4 + POC, and the total BC was defined as EC1 + EC2 + EC3-POC. Char was defined as EC-POC, and soot was defined as EC2 + EC3, as defined by Han et al [34].

#### 2.3.2 LFBC and HFBC measurement

An improved version of the methodology used by Janzen et al [35] was adopted. In brief, the light group organic matter suspended on the surface of the BC mixture was separated using the dichromate oxidation method by using a hydrofluoric/hydrochloric acid treatment, the supernatant was vacuum filtered through a 0.45 μm filter membrane, and the resulting sample was washed with 100 ml 0.01 mol·L^-1^CaCl_2_ solution and repeatedly rinsed with 200 ml distilled water to obtain light-fraction black carbon (LFBC); the precipitate in the aforementioned centrifuge tube was added with 50 ml distilled water, shaken for 0.5 h (200 r·min^-1^), and centrifuged for 20 min at 4000 r·min^-1^; the supernatant was discarded, and washing was repeated three times; and the precipitate in the tube was repeatedly washed with 95% ethanol until colorless and was divided into heavy-fraction black carbon (HFBC).

#### 2.3.3 Soils physic-chemical properties

Soil total OC (TOC), OC, and total nitrogen (TN) were determined using a Vario ELIII elemental analyzer (Elementar GMBH). Total phosphorus (TP) was determined using concentrated sulfuric acid-perchloric acid digestion and the molybdenum antimony anti-colorimetric method (Agilent 8453 UV-Vis spectrophotometer, USA). Total potassium (TK) was determined using sulfuric acid-perchloric acid digestion and the flame photometric method. Fast-acting nitrogen (AN) was determined using the alkaline digestion diffusion method. Fast-acting phosphorus (AP) was determined using sodium bicarbonate leaching and the molybdenum antimony anti-colorimetric method. Fast-acting potassium (AK) was determined using the flame photometric method. Fresh weight was determined using the ring knife method and returned to the laboratory for drying and weighing to calculate the soil water content and bulk weight (BD) [36].

### 2.4 Statistical analytics

The experimental data were tabulated using Excel (2007). SPSS (19.0) was used for significance testing and correlation analysis. Data analysis and graphing were performed using Origin the 2019 software.

## 3 Results

### 3.1 Variations in physical and chemical properties in soils of different land uses

Physical and chemical properties in soils of different land uses are presented in Fig 2. The TN (Fig 2a) values of secondary forest profiles (1.96–4.20 g·kg^-1^) were significantly higher than those of shrub profiles (1.18–2.50 g·kg^-1^), farmland profiles (0.99–2.18 g·kg^-1^), and wasteland profiles (0.97–1.82 g·kg^-1^) in each soil layer. The TP (Fig 2b), TK (Fig 2c), AN (Fig 2d), AP (Fig 2e), AK (Fig 2f), and TOC (Fig 2g) values in secondary forest profiles were significantly higher than those in shrub, farmland, and wasteland profiles in each soil layer. The BD (Fig 2h) values of secondary forest profiles (1.07–1.38 g·cm^-3^), farmland profiles (1.18–1.48 g·cm^-3^), and wasteland profiles (1.25–1.50 g·cm^-3^) were lower than those of wasteland profiles (1.37–1.52 g·cm^-3^). TN, TP, TK, AN, AP, AK, and TOC in the soils of secondary forest, shrub, farmland, and wasteland fields tended to decrease with increasing depth, and BD tended to increase with depth. TN, TP, TK, AN, AP, AK, TOC, and BD significantly differed among the different layers of the same land use.

**Fig 2.**
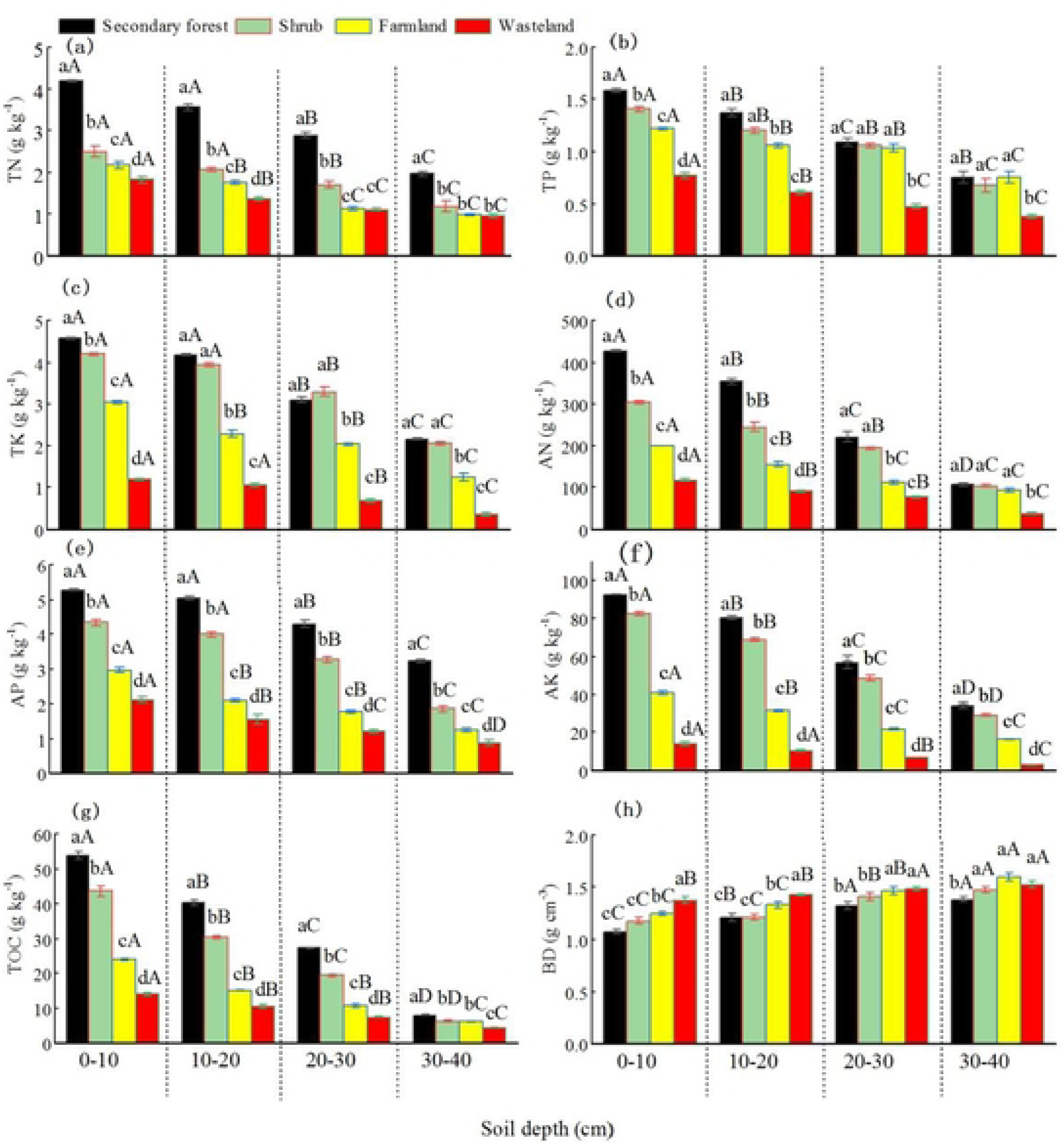
Physical and chemical properties in soils of different land uses. Note: Lower case letters indicate significant differences (*P*<0.05) between different land use types in the same layer, while upper case letters indicate significant differences (*P* <0.05) between different layers in the same land use type. The same as follows.

### 3.2 Characteristics of changes in SOC and BC content of different land uses

The OC, BC, char, and HFBC contents tended to decrease with increasing depth across the four land use patterns (Fig 3). Soot content tended to decrease with increasing depth in secondary forest, farmlands, and wastelands, and LFBC content decreased with increasing depth in secondary forest and shrub. The δ^13^C_BC_ and BC/TOC ratio increased with increasing depth for the four land use patterns. The OC (3.14–42.78 g·kg^-1^), BC (1.95–9.70 g·kg^-1^), and HFBC (1.12–6.09 g·kg^-1^) values of secondary forest were higher than those of shrub, farmland, and wasteland in each soil layer. There were no significant differences in OC content between farmland and wasteland in the 20–30 cm layer and among shrub, farmland, and wasteland in the 30–40 cm layer. The δ^13^C_BC_ content in the 0–20 cm layer showed a significant difference among the four land use methods, with wasteland having the highest value, followed by farmland, shrub, and secondary forest (*P<*0.05); the values of δ^13^C_BC_ had significant differences between secondary forest and other land use patterns in the 30– 40 cm layer.

**Fig 3.**
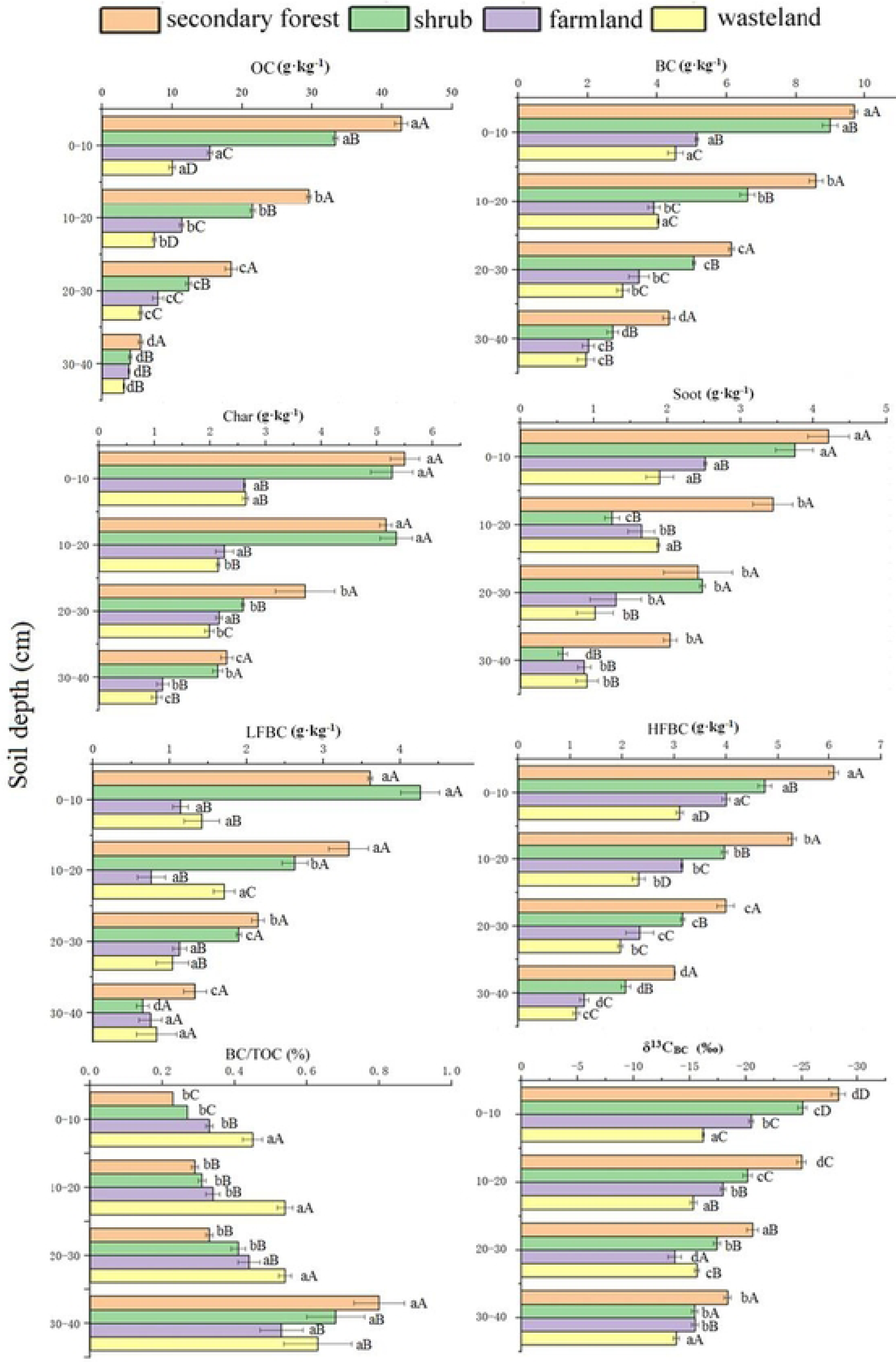
Characteristics of OC, BC, char, and soot concentrations and BC/TOC ratios in soil under different laud uses.

Char content ranged from 1.04 to 5.50 g·kg^-1^ in each soil layer. The char values of secondary forest were higher than those of shrub, wasteland, and farmland in the 0– 10 cm layer and in the 20–40 cm layer. The char values of shrub were higher than those of secondary forest, farmland, and wasteland in the 10–20 cm layer. Soot content ranged from 0.58 to 4.21 g·kg^-1^ across each soil layer. The values of secondary forest were significantly higher than those of shrub, wasteland, and farmland in the 0–10, 10–20, and 30–40 cm layers; nevertheless, the soot values of shrub were significantly higher than those of secondary forest, wasteland, and farmland in the 20–30 cm layer.

The LFBC content of the four land uses varied from 0.65 to 4.26 g·kg^-1^. The LFBC values of shrub were higher than those of secondary forest, wasteland, and farmland in the 0–10 cm layer. The LFBC values of secondary forest were higher than those of shrub, wasteland, and farmland in the 10–40 cm layer. The BC/TOC values of wasteland were significantly higher than those of secondary forest, shrub, and farmland in the 0–30 cm layer. In general, the OC, BC, char, soot, LFBC and HFBC concentrations of secondary forest and shrub were significantly higher than those of farmland and wasteland in the 0–40 cm layer.

### 3.3 Effects of various land use patterns on proportion of soil BC composition

Soil BC composition ratios are shown in Table 1. The BC/OC values of the four land use patterns were secondary forest (22.67%–78.62%), shrub (27.02%–67.16%), farmland (33.29%–52.47%), and wasteland (45.16%–62.10%) in the 0–40 cm layer, respectively. BC/OC tended to increase with depth, reaching its highest level in the 30–40 cm layer, and the BC/OC values of secondary forest (78.62%) were significantly higher than those of shrub (67.16%), wasteland (62.10%), and farmland (52.47%). Char/BC varied irregularly with soil depth, and the char/BC values of shrub were higher than those of secondary forest, wasteland, and farmland in the 0–10, 10– 20, and 30–40 cm layers. The char/BC values of shrub were lower than those of secondary forest, wasteland, and farmland in the 20–30 cm layer. Influenced by land use, char/soot exhibited a pattern of change similar to that of char/BC. Soot/BC changed unpredictably, and each layer showed different characteristics. Soot/BC values of farmland were higher than those of secondary forest, wasteland, and shrub in the 0–10 cm layer. In the 10–20 cm layer, the values of wasteland were higher than those of farmland, secondary forest, and shrub. The soot/BC values of secondary forest were higher than those of wasteland, farmland, and shrub in the 30–40 cm layer.

The LFBC/BC values of shrub were significantly higher than those of secondary forest, wasteland, and farmland in the 0–10 and 20–30 cm layers. The LFBC/BC values of wasteland were significantly higher than those of secondary forest, shrub, and farmland in the 10–20 and 30–40 cm layers. The HFBC/BC values of farmland were significantly higher than those of secondary forest, wasteland, and shrub in the 0–10, 10–20, and 20–30 cm layers, and the HFBC/BC values of shrub were significantly higher than those of secondary forest, wasteland, and farmland in the 30–40 cm layer. HFBC/BC was higher than LFBC/BC in different soil use patterns, indicating a high contribution of heavy-fraction black carbon to soil black carbon.

### 3.4 Correlation between soil BC and soil physicochemical properties

The correlation between soil BC fraction content and soil physicochemical properties in the 0–40 cm soil layer was analyzed Fig 4 shows that OC, TN, TP, TK, AN, AP, AK, and SOC were highly significantly positively correlated with BC, char, soot, LFBC, and HFBC (*P<0*.001); OC and SOC were significantly positively correlated with LFBC/BC (*P<0*.05); TN, TP, TK, AN, AP, AK, and BD were not correlated with LFBC/BC and HFBC/BC; OC and SOC were significantly and negatively correlated with HFBC/BC (*P*<0.05); BD was highly significantly and negatively correlated with BC, char, soot, LFBC, and HFBC (*P*<0.001) and was highly and significantly and positively correlated with δ^13^C_BC_ (*P*<0.001); and OC, TN, TP, TK, AN, AP, AK, and SOC were highly significantly negatively correlated with BC/OC (*P*<0.001). Accordingly, there was a strong correlation between the BC content of BC components in the soil and the physical and chemical properties of the soil.

**Fig 4.**
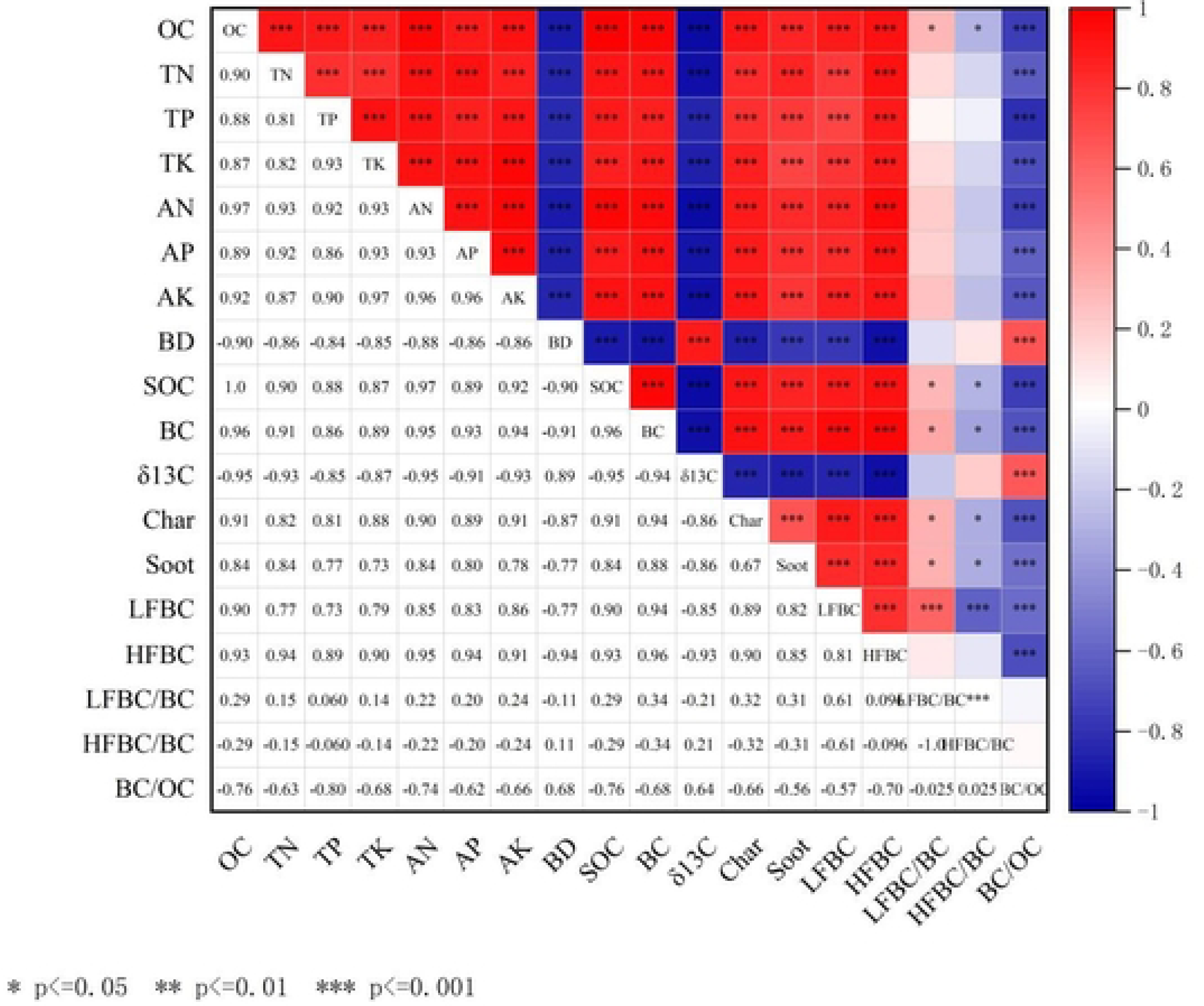
Correlation between soil BC fraction and soil physicochemical properties.

The first (RD1) and second (RD2) standard axes of the soil BC fraction accounted for 69.38% and 7.91% of the variation in the soil BC fraction, respectively (Fig 5). Correlation analysis revealed that BC was significantly and positively associated with the OC content, char, soot, LFBC, TK, AP and HFBC. Additionally, there were highly significant and positive correlations between BC and soil δ^13^C_BC_ and HFBC/BC (*P<*0.05). OC had highly significant negative correlations with soil BC stock, BD and LFBC/BC. It showed a positive correlation among OC, LFBC and HFBC contents. It also revealed that OC and component analysis were inversely related. Furthermore, the correlation between LFBC and BC was weaker than that between HFBC and BC, suggesting that the heavy fraction BC had a stronger association with OC than the light fraction BC.

**Fig 5.**
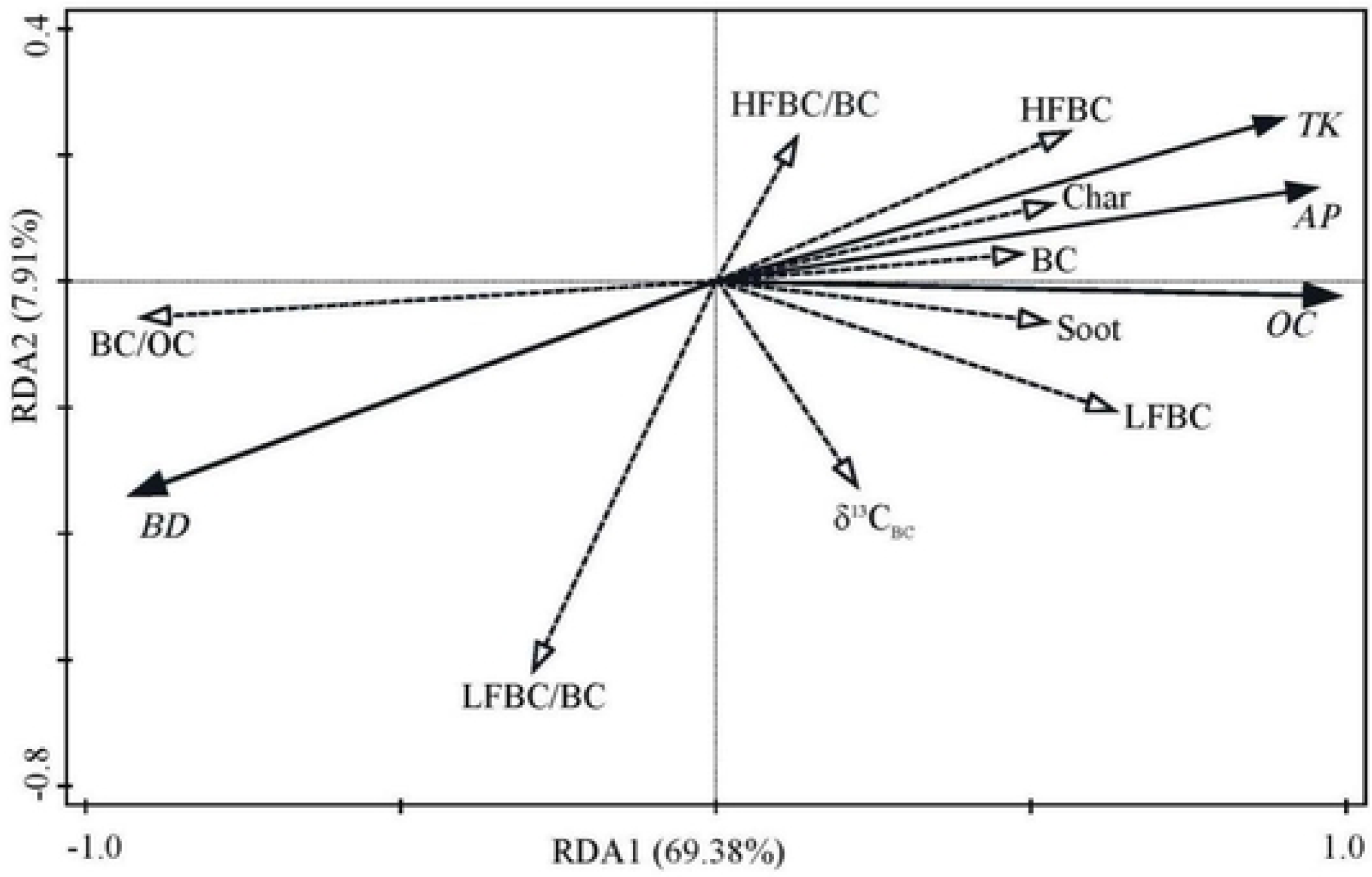
Redundancy analysis of soil BC fraction and soil physicochemical properties.

## 4 Discussion

### 4.1 Characteristics of soil BC in rocky desertification in Southwest Guangxi

BC, as a marker of human activities, is widely present in soils affected by anthropogenic disturbances [37]. The average BC content in soils from rock-deserted areas in Southwest Guangxi ranged from 3.38 to 7.20 g·kg^-1^, which is comparable to the natural soil BC content of (2 ± 1)–(6 ± 3) g·kg^-1^ observed by Karthik et al [38]in South India; higher than the BC content of 2.99–4.36 g·kg^-1^ in woodland soil observed by Jiang et al [39]; lower than that of 6.39–16.55 g·kg^-1^ for Changbai Mountain, China, observed by Sun et al [40]; and lower than that of 6.64–17.63 g·kg^-1^ in the northeastern forest area observed by Sun et al [41]. The low BC content of the soils of the study area is primarily attributed to the unique habitats and climatic conditions of the rocky desertification region. In rocky desertification areas, the soil formation process is slow, the soil layer is shallow, soil leakage is strong, and the soil lacks weathered parent material, causing soil erosion and irreversible damage [42].

Land use is an important controlling factor for SOC fractions [16] and in this study, land use patterns had a significant impact on the BC content and proportion to TOC. The mean BC content at the 0–40 cm depth varied among the four land use patterns—secondary forest (7.20 g·kg^-1^), shrub (5.85 g·kg^-1^), farmland (3.64 g·kg^-1^), and wasteland (3.38 g·kg^-1^)—indicating that the multi-year measures of vegetation restoration had contributed to the increase in soil BC content. The highest BC concentration was in the surface layer, with a gradual decrease as depth increased. These findings are consistent with those of Rumpel et al [43] and Sun et al [40]. There was some variation in the proportion of soil BC to TOC among the land use patterns within the same layer. The mean BC content was highest in wasteland (32.24%), followed by farmland (21.44%), shrub (20.73%), and secondary forest (18.06%) in the 0–10 cm soil layer. There was also a significant difference between soil BC and the proportion of TOC within different soil layers under the same land use.

The rocky desertification area in Southwest Guangxi has a long history of anthropogenic farming, resulting in severe rocky desertification and a degraded ecological environment due to human–land conflict [42]. Benefitting from the successful implementation of the national policy called the Grain-for-Green project, the local ecology of rocky desertification areas has been effectively restored to a certain extent [42]. With the restoration of vegetation, secondary forest and shrub accumulated SOC, increasing the high soil BC content. The return and decomposition of significant amounts of organic matter, such as dead branches and leaves, further contributes to the maintenance of soil carbon content, enhancing its nutrient quality and promoting the maintenance of soil carbon, leading to an increase in soil nutrient content and enhancement of OC levels. This improved soil structure provides the secondary forest and shrub areas with ample nutrient sources, suitable moisture levels, and optimal aeration. These conditions create favorable environments for increasing soil microorganism activity and organic matter decomposition.

In terrestrial ecosystems, the SOC content is controlled by complex interactions among C inputs, C stabilization processes, and C loss from the soil [16]. By contrast, existing farmland was treated solely with chemical fertilizers for an extended period. Relatively loose and bare soil surfaces destroyed the soil aggregates. Although farmyard manure application has the potential to increase soil BC content, it remains underutilized [44].

The flushing of abundant rainfall has resulted in significant soil carbon loss, exacerbated by human production activities that predominantly adopt a material output-based farming approach. Furthermore, the majority of the organic matter generated by crops is removed from the system, leading to reduced nutrient cycling and soil fertility, which are not conducive to SOC, microbial load, and activity. The exposed surface area of wasteland resulted in a decrease in SOC due to severe erosion, leading to a significant reduction in microbial quantity and activity. In this study, the contents of OC, BC, char, soot, LFBC, and HFBC were lower in each layer in the 0– 40 cm depth in farmland and wasteland; the lowest levels were in wasteland, which could be attributed to anthropogenic tillage practices, fertilization activities, and loss of organic matter from the surface that damaged the soil structure and impeded the return of organic matter to the soil.

The soil BC content was significantly higher in secondary forest and shrub than in farmland and wasteland, owing to varying degrees of soil carbon input and differences in vegetation effects on SOC. Additionally, anthropogenic disturbance measures such as fire have historically existed in secondary forest and shrub areas, increasing the regulation of BC within the soil.

### 4.2 Correlation between BC and OC in the rocky desertification region of Southwest Guangxi

In this study, a significant positive correlation was observed between soil BC and OC, indicating that BC plays an important role in SOC fixation. These findings are consistent with those of existing studies [33, 41, 45], suggesting that an increase in BC content of secondary forest and shrub areas contributes to SOC accumulation. On the one hand, the SOC accumulation is attributed to the unique chemical and biological inertness of BC; on the other hand, BC can adsorb and fix organic matter and clay minerals. Furthermore, BC and OC exhibited a surface enrichment pattern and decreased significantly with depth in the soil profile, which is consistent with the findings of Sun et al [40].

The ratio of soil BC to OC under the four different land use patterns increased proportionally with increasing soil depth. Possible reasons for this are as follows: ① A significant amount of litter in secondary forest and shrub is annually enriched in the surface soil, promoting OC accumulation. OC naturally acts as a “diluent” for BC without human interference. Although frequent disturbances occur on farmland surfaces, the application of large amounts of organic matter decreases fresh carbon input; however, the effect of carbon on soil depth is not apparent. Different land uses can strengthen the degradation capacity of newly imported carbon sources and BC because microbial activity in the upper layer of soil with abundant oxygen is high. Conversely, the lower soil layer has low microbial activity and poor permeability, which reduces the possibility of BC degradation [45]. ③ BC is highly resistant to chemical inert and thermal stability [37]. When migrating downward in the soil, BC and non-BC components exhibited different levels of stability in the lower layers. As a result, BC is selectively enriched in the soil during long-term soil formation [46]. The migration rates of BC and non-BC components may differ; BC has higher migration rates than non-BC components.

In this study, the percentage of BC in heavy fractions was significantly higher than that in light fractions, which contradicted the results reported by Glaser et al [47], who found that BC mainly existed in light fractions of Amazonian black soils. Brodowski et al [44] found that BC in German soils was predominantly present in light fractions due to the inert nature of HFBC and its inclusion within the protective fraction of soil carbon, which is associated with the unique geological background environment of karstic rocky desertification areas. Brodowski et al [44] also wasdemonstrated that due to the high chemical and thermal stability of BC, its chemical composition and properties are similar to those of highly aromatic soil humic acid. Moreover, the proportion of highly aromatic carbon increases with soil depth [48], indicating that BC decreases less significantly with increasing soil depth than TOC. Under similar conditions of climate, soil-forming parent material, and topography, litter serves as the primary supplement to SOC, and the influx of substrate carbon into the system significantly affects the increase in BC [49]. The amount of desiccated twigs and decayed foliage entering the soil is determined by aboveground vegetation, and decomposition rates vary among plants. Different land use patterns result in varying types, yields, and quality of litter [50], and the levels of BC input into the soil vary, indirectly influencing the microenvironment of soil microbial activity [51].

### 4.3 Influence of land uses on soil BC composition

The proportions of BC to OC in this study were 22.67%–78.62% for secondary forest, 27.02%–67.16% for shrub, 33.29%–52.47% for farmland, and 45.16%–62.10% for wasteland (Table 1), higher than those in existing studies, such as the 8%–26% in Zhang et al [52] in red and yellow soil, and 11.3%–53.2% in Dai et al [53] in agricultural soil in the northern Zhejiang Plain. This may be attributed to the low OC content of rocky desertification areas. Additionally, the distribution of BC/OC in the surface soil was the highest in wasteland (45.16%), followed by farmland (33.29%), scrub (27.02%), and secondary forest (22.67%). These results are consistent with those of Zhang et al [52], who found the highest OC content for dry land, followed by tea gardens, secondary forest, and native woodland, which may be due to a gradual decrease in OC content from secondary forest to cultivated soils with increasing tillage time. Additionally, BC exhibits high stability, and organic fertilizers often contain incompletely burned biochar. Consequently, the BC/OC ratio tended to be high in the soil layers with the application of organic fertilizers, suggesting that agricultural activities influence the accumulation of soil BC.

In this study, BC showed a significant positive correlation with char and soot content. The char/BC ratio varied from 51.09% to 81.01%, and the soot/BC ratio varied from 18.99% to 49.09%. These results indicate that char and soot are important components of soil BC in the rocky desertification areas of Southwest Guangxi, similar to the findings of Zhan et al [33]. The BC/TOC ranged from 18.06% to 54.41% (Fig 3), indicating significant variations and the importance of BC as a major component of SOC, with much higher content than that of Zhang et al [52] (8.3%– 25.6%). The decomposition rate of the original OC in the soil is influenced by the amount of BC input [54], and the high BC/TOC observed in this study may be attributed to either a great input of BC or low levels of OC within the soil. Different land uses had distinct impacts on BC/TOC, with wasteland having the highest ratio (39.1%), followed by secondary forest (29.10%), farmland (28.22%), and shrub (28.16%). Thus, BC is additional influenced by human activities. Agricultural tillage can lead to a reduction in TOC content, whereas the application of organic fertilizers and plant litter input can increase surface SOC content.

This study found that soil bulk density was negatively correlated with BC. The soil bulk density gradually decreased in the order wasteland, farmland, shrub, and secondary forest, and its size reflected the soil structure and water retention capacity. The permeability of the surface soil was affected by artificial tillage; therefore, its bulk density was low. However, the bulk density of wastelands is larg because of human disturbance and a lack of effective management measures. Secondary forest and shrub areas have favorable environmental conditions, where stable, healthy natural systems form and, therefore, have small bulk density values. This phenomenon might also explain the differences in BC/TOC, consistent with the results of Zhan et al [5] and Edmondson et al [55].

BC is mainly generated from two sources: natural emissions, primarily resulting from volcanic eruptions and fires in grasslands or forests, and anthropogenic emissions, primarily resulting from residential and commercial activities, energy production, industrial processes, motor vehicle exhaust, coal combustion, and agricultural straw burning. When analyzing the sources of BC in soil, many scholars [5, 56–57] have used the char/soot method. Generally, char is formed by the incomplete combustion of biomass, and soot is predominantly produced by fossil fuel combustion. Therefore, BC source analysis using char/soot has a substantially reference value. Zhan et al [5] reported a char/soot ratio of less than 2.0, suggesting that the primary sources of BC are herbaceous plants and fossil fuel combustion, such as motor vehicle exhaust emissions. Conversely, a ratio greater than 2.0 indicates that woody plant combustion is the main contributor to BC. This study indicated that the char/soot ratio was below 2.0 across all layers (Table 1), except 4.38 for the 10–20 cm depth, 3.73 for the 30–40 cm depth in shrub, and 2.34 for the 20–30 cm depth of wasteland, indicating that herbaceous plants and fossil fuel combustion are the primary sources of soil BC within the study area. This was mainly related to the succession sequence of herbs, shrubs, and trees during the process of vegetation restoration. Another contributing factor might be that the study area is close to roads, dust, and vehicle exhaust emissions from road traffic. Stable carbon isotope values from -13.66‰ to -28.33‰ and the high proportion of HFBC in the soil BC suggest that the BC is mainly from the burning of C3 plants [58], as well as motor vehicle exhaust emissions and fossil fuel combustion [37].

The isotopic composition of BC from C3 and C4 plants is relatively fixed. Das et al [59] showed that the δ^13^C PDB values of BC were approximately from -24. 6‰ to -26.1‰ for C3 plants and from -12.3‰ to -13.8‰ for C4 plants after being burned. When a substantial quantity of soil BC particles results from the combustion of C3 plants, such as that generated by forest fires and wood burning, its δ^13^C value gradually becomes BC originating from C3 plants with the increasing amount of BC. Due to the extensive combustion of fossil fuels, the BC particles exhibited a relative dilution of δ^13^C, leading to a reduction in atmospheric CO_2_ δ^13^C values [60] and the δ^13^C value of BC of carbon-contained materials on the soil surface layer also decreased. The BC content of secondary forest soils was significantly higher than that of other land use types, and the ^13^C abundance of soil BC was the lowest, indicating a great depletion of ^13^C in BC. In secondary forest, the many years of burning biomass materials and road motor vehicle exhaust containing large amounts of particulate matter, such as BC, resulted in a high soil BC content. The δ^13^BC values were lower than those in other land use modalities, indicating that secondary forest and shrub reduced the diffusion of BC from motor vehicle emissions to surrounding areas. Road traffic dust may also significantly contribute to soil BC in the study area because the amount of BC accumulated in road dust is primarily derived from fossil fuel combustion emissions from motor vehicles [34]. This suggests that vehicular emission pollution has become a potential source of air pollution with the development of tourism and the increased number of motor vehicles in the rocky desertification areas of Southwest Guangxi.

## 5 Conclusions

The BC content exhibited a significant positive correlation with OC. The mean BC content of the 0–40 cm soil layer was the highest in secondary forest, followed by shrub, farmland, and wasteland. Land use significantly influenced the soil BC distribution, with a decreasing trend observed after conversion from forest land to farmland. The enhancement of vegetation restoration measures and the reduction of anthropogenic reclamation are therefore recommended. The contribution of BC to SOC was positively correlated with soil depth, and the increase in soil BC content led to the accumulation of soil. The proportion of BC in the heavy fraction OC was significantly higher than that in the light fraction OC. Soil char, soot, and light and heavy BC were positively correlated with BC content. BC improved the carbon sequestration capacity of secondary forest soils in the rocky desertification area of Southwest Guangxi, and vegetation restoration significantly contributed to BC accumulation. Land use and human activities are the main factors affecting soil black carbon distribution in this area.

## Author contributions

**Conceptualization:** Denan Zhang, Qiumei Teng, Kechao Huang, Guangping Xu.

**Data curation:** Denan Zhang, Qiumei Teng, Kechao Huang.

**Formal analysis:** Yuyi Shen, Yingjie Sun.

**Funding acquisition:** Denan Zhang, Qiumei Teng, Guangping Xu. **Investigation:** Denan Zhang, Qiumei Teng, Shihong Lyu,Yanzhao Zhang. **Supervision:** Shihong Lyu,Yanzhao Zhang.

**Visualization:** Denan Zhang.

**Writing (original draft):** Denan Zhang.

**Writing (review and editing):**Denan Zhang, Guangping Xu.

All authors have read and agreed to the published version of the manuscript.

## Funding

This work was financially supported by the National Natural Science Foundation of China(42267007;32460311);Natural Science Foundation of Guangxi (2020GXNSFBA297048); Basic Research Fund of Guangxi Academy of Sciences (CQZ-E-1912); Guangxi Key Science and Technology Innovation Base on Karst Dynamics (KDL & Guangxi 202004); Fund of Guangxi Key Laboratory of Plant Conservation and Restoration Ecology in Karst Terrain (22-035-26); Basic Research Fund of Guangxi Institute of Botany (23006; 22003).

## Institutional Review Board Statement

Not applicable.

## Data Availability Statement

The data presented in this study are available on request from the corresponding author.

## Conflicts of Interest

The authors declare no conflicts of interest.

## Acknowledgments

We would like to thank Jianchun Liu, Guixia Cheng, Qianqian Yu, Lei Tian, Qilei Cheng, and Cuihong Li for testing the server.

## Notes

### Competing Interest Statement

The authors have declared no competing interest.

## References

1. Ramesh, T., Bolan, N.S., Kirkham, M.B., Wijesekara, H., Kanchikerimath, M., Rao, C.S., & Freeman II, O.W. Soil organic carbon dynamics: Impact of land use changes and management practices: A review. Advances in agronomy. 2019,156, 1–107. 10.1016/bs.agron.2019.02.001

2. Schmidt, M.W., &Noack, A.G. Black carbon in soils and sediments: analysis, distribution, implications, and current challenges. Global biogeochemical cycles.2000,143, 777–793. 10.1029/1999GB001208

3. Forbes, M.S., Raison, R.J.,& Skjemstad, J.O. Formation, transformation and transport of black carbon charcoal in terrestrial and aquatic ecosystems. Science of the total environment. 2006, 3701, 190–206. 10.1016/j.scitotenv.2006.06.007

4. Glaser, B., & Birk, J.J. State of the scientific knowledge on properties and genesis of Anthropogenic Dark Earths in Central Amazonia terra preta de Índio. Geochimica et Cosmochimica acta, 2012, 82, 39–51. 10.1016/j.gca.2010.11.029

5. Zhan, C.L., Cao, J.J., Han, Y.M., Wang, P., Huang, R.J., Wei, C., Hu, W.G., &Zhang, J.Q. Spatial patterns, storages and sources of black carbon in soils from the catchment of Qinghai Lake, China. European Journal of Soil Science. 2015, 663, 525–534. 10.1111/ejss.12236

6. Gao, H., Li, H.X., Shi, J.Q., Huang, J.B., Wei, J., Qu, X.L., &Long, T. Black carbon, soil organic matter molecular signatures under different land uses in Shenyang. China and relationship with PAHs, Chemosphere. 2023, 342, 140089. 10.1016/j.chemosphere.2023.140089

7. Sharma, R., Mishra, A.K., Role of essential climate variables and black carbon in climate change: Possible mitigation strategies. Biomass, Biofuels, Biochemicals. 2022, 31–53. 10.1016/B978-0-12-823500-3.00005-4

8. Preston, C.M., &Schmidt, M.W. Black pyrogenic carbon: a synthesis of current knowledge and uncertainties with special consideration of boreal regions. Biogeosciences. 2006, 34, 397–420. 10.5194/bg-3-397-2006

9. Lehmann, J. A handful of carbon. Nature. 2007, 4477141, 143–144. 10.1038/447143a

10. Biederman, L.A., &Harpole, W.S. Biochar and its effects on plant productivity and nutrient cycling: a meta-analysis. GCB bioenergy. 2013, 52, 202–214. 10.1111/gcbb.12037

11. Ohlson, M., Dahlberg, B., Økland, T., Brown, K.J., &Halvorsen, R. The charcoal carbon pool in boreal forest soils. Nature Geoscience. 2009, 210, 692–695. 10.1038/ngeo617

12. Lorenz, K., Lal, R., &Jiménez, J.J. Characterization of soil organic matter and black carbon in dry tropical forests of Costa Rica. Geoderma. 2010, 1583–4, 315-321. 10.1016/j.geoderma.2010.05.011

13. Zhang, T.T., Wooster, M.J., Green, D.C., &Main, B. New field-based agricultural biomass burning trace gas, PM2.5, and black carbon emission ratios and factors measured in situ at crop residue fires in Eastern China. Atmospheric Environment. 2015, 121, 22-34. 10.1016/j.atmosenv.2015.05.010

14. Wang, Q., Zhang, P.J., Liu, M., and Deng, Z.W. Mineral-associated organic carbon and black carbon in restored wetlands. Soil Biology and Biochemistry. 2014a, 75, 300–309. 10.1016/j.soilbio.2014.04.025

15. Ji, D.S., Li, L., Pang, B., Xue, P., Wang, L.L., Wu, Y.F., Zhang, H.L., &Wang, Y.S. Characterization of black carbon in an urban-rural fringe area of Beijing. Environmental Pollution. 2017, 223, 524–534. 10.1016/j.envpol.2017.01.055

16. Qin, Z.L., Yang, X.M., Song, Z.L., Peng, B., Van Zwieten, L., Yu, C.X., Wu, S.C., Mohammad, M., & Wang, H.L. Vertical distributions of organic carbon fractions under paddy and forest soils derived from black shales: Implications for potential of long-term carbon storage. Catena. 2021, 198, 105056. 10.1016/j.catena.2020.105056

17. Gao, G.Y., Tuo, D.F., Han, X.Y., Jiao, L., Li, J., &Fu, B.J. Effects of land-use patterns on soil carbon and nitrogen variations along revegetated hillslopes in the Chinese Loess Plateau. Science of the Total Environment. 2020, 746, 141156. 10.1016/j.scitotenv.2020.141156

18. Valjavec, M.B., Čarni, A., Žlindra, D., Zorn, M., &Marinšek, A. Soil organic carbon stock capacity in karst dolines under different land uses. Catena. 2022, 218, 106548. 10.1016/j.catena.2022.106548

19. Davari, M., Gholami, L., Nabiollahi, K., Homaee, M., &Jafari, H.J. Deforestation and cultivation of sparse forest impacts on soil quality case study: West Iran, Baneh. Soil and Tillage Research. 2020, 198, 104504. 10.1016/j.still.2019.104504

20. Peng, X.Y., Huang, Y., Duan, X.W., Yang, H., &Liu, J.X. Particulate and mineral-associated organic carbon fractions reveal the roles of soil aggregates under different land-use types in a karst faulted basin of China. Catena. 2023, 220, 106721. 10.1016/j.catena.2022.106721

21. Jiang, Z.C., Luo, W.Q., Tong, L.Q., Cheng, Y., Yang, Q.Y., Wu, Z.Y., &Liang, J.H. Evolution features of rocky desertification and influence factors in karst areas of southwest China in the 21st century. Carsologica Sinica. 2016, 355, 461–468. 10.11932/karst20160504

22. Uchida, E., Xu, J.T., &Rozelle, S. Grain for green: Cost-effectiveness and sustainability of China’s conservation set-aside program. Land Economics. 2005, 812, 247–264. http://about.jstor.org/terms

23. Xu, G.P., He, C.X., Zhang, D.N., Zhao, Z.G., Lu, S.H., Yao, Y.F., &Huang, Y.Q. Soil microbial biomass and active characters under different land-use types in karst mountain areas of southwest Guangxi. Guihaia. 2013, 333, 331–337. 10.3969/j.issn.1000-3142.2013.03.009

24. Lyu, S.H., Li, X.Q., Bai, K.D., Pan, Y.M., Tang, S.C., Deng, L.L., &Zeng, D.J. Microtopographic differentiation characteristics of soil physicochemical properties and leaf traits of Litsea glutinosa in karst rocky desertification mountain of southwestern Guangxi. Journal of Plant Resources and Environment. 2022, 313, 11–17. 10.3969/j.issn.1674-7895.2022.03.02

25. Kang, Z.Q., Chen, J., Yuan, D.X., He, S.Y., Li, Y.L, Chang, Y., Deng, Y., Chen, Y., Liu, Y.Y., Jiang, G.H., Wang, X.Y., &Zhang, Q. Promotion function of forest vegetation on the water & carbon coupling cycle in karst critical zone: Insights from karst groundwater systems in south China. Journal of Hydrology. 2020, 590, 125246. 10.1016/j.jhydrol.2020.125246

26. Mo, C.X., Lai, S.F., Yang, Q., Huang, K.K., Lei, X.B., Yang, L.F., Yan, Z.W., &Jiang, C. A comprehensive assessment of runoff dynamics in response to climate change and human activities in a typical karst watershed, southwest China. Journal of Environmental Management. 2023, 332, 117380. 10.1016/j.jenvman.2023.117380

27. Xu, G.P., Gu, D.X., Pan, F.J., Su, Y.J., Luo, A.Y., He, C.X., &Huang, Y.Q. Effects of different land-use types on soil enzyme activity in karst mountain areas of Southwest Guangxi. Guihaia. 2014, 4,460–466. 10.3969/j.issn.1000-3142.2014.04.006

28. Qiu, S.J., Peng, J., Zheng, H.N., Xu, Z.H., &Meersmans, J. How can massive ecological restoration programs interplay with social-ecological systems? A review of research in the South China karst region. Science of The Total Environment. 2022, 807, 150723. 10.1016/j.scitotenv.2021.150723

29. He, H.Z., Zhang, Z.M., Liu, Y.Y., &Sun, C. Distribution Characteristics of Organic Carbon and Black Carbon in Karst Forest Soil of Yuntai Mountain of Guizhou Province. Guizhou Agricultural Sciences. 2013, 415,90–92. 10.3969/j.issn.1001-3601.2013.05.027

30. Nobuhisa K, Seiji S, Yasuhito S, Takashi K, Takeo S, Hiroshi N, et al. Assessing changes in soil carbon stocks after land use conversion from forest land to agricultural land in Japan. Geoderma. 377, 2020, 114487. 10.1016/j.geoderma.2020.114487

31. Lyu, S.H., Li, X.Q., Bai, K.D., Wei, C.Q., Zeng, D.J., &Xu, G.P. The effects of three pioneer tree species on facilitation and twig and leaf traits of Cyclobalanopsis glauca seedlings in a rocky desertification region of Guangxi, China. Chinese Journal of Ecology. 2018, 377,1917–1924. 10.13292/j.1000-4890.201807.024

32. Lim, B., &Cachier, H. Determination of black carbon by chemical oxidation and thermal treatment in recent marine and lake sediments and Cretaceous-Tertiary clays. Chemical Geology. 1996, 1311-4, 143–154. 10.1016/0009-2541(96)00031-9

33. Zhan, C.L., Cao, J.J., Han, Y.M., Huang, S.P., Tu, X.M., Wang, P., &An, Z.S. Spatial distributions and sequestrations of organic carbon and black carbon in soils from the Chinese loess plateau. Science of the Total Environment. 2013, 465, 255–266. 10.1016/j.scitotenv.2012.10.113

34. Han, Y.M., Cao, J.J., Chow, J.C., Watson, J.G, An, Z.S., &Liu, S.X. Elemental carbon in urban soils and road dusts in Xi’an, China and its implication for air pollution. Atmospheric Environment. 2009, 4315, 2464–2470. 10.1016/j.atmosenv.2009.01.040

35. Janzen, H.H., Campbell, C.A., Brandt, S.A., Lafond, G.P., &Townley-Smith, L. Light-fraction organic matter in soils from long-term crop rotations. Soil Science Society of America Journal. 1992, 566, 1799–1806. 10.2136/sssaj1992.03615995005600060025x

36. Bao, S. Soil Agrochemical analysis. Beijing: China Agriculture Press, in Chinese. 2000.

37. Koç, K., Koşun, E., Cheng, H., Demirtaş, F., Edwards, R.L., &Fleitmann, D. Black carbon traces of human activities in stalagmites from Turkey. Journal of Archaeological Science. 2020, 123, 105255. 10.1016/j.jas.2020.105255

38. Karthik, V., Bhaskar, B.V., Ramachandran, S., &Kumar, P. Black carbon flux in terrestrial and aquatic environments of Kodaikanal in the Western Ghats, South India: Estimation, source identification, and implication. Science of The Total Environment. 2023, 854, 158647. 10.1016/j.scitotenv.2022.158647

39. Jiang, D., He, Z.M., Yin, Y.F., Fan, S.H., Huang, Z.Q., &Wan, X. H. Effects of Harvest Residue Management on Black Carbon and Black Nitrogen in Plantation Soil. Journal of Subtropical Resources and Environment. 2014a, 3, 68–74. 10.3969/j.issn.1673-7105.2014.03.009

40. Sun, J.B., Sang, Y., Song, J.F., &Chui, X.Y. Content and distribution of black carbon in typical forest soils in changbaishan mountains. Forest Research. 2016, 291, 34–40. 10.3969/j.issn.1001-1498.2016.01.005

41. Sun, J.B., Xu, J.H., Song, J.F., &Chui, X.Y. Distribution characteristics and influence factors of soil black carbon of typical forest areas in northeast China. Acta Scientiae Circumstantiae. 2018, 388, 3313–3321. 10.13671/j.hjkxxb.2018.0172

42. Jiang, Z.C., Lian, Y.Q., &Qin, X.Q. Rocky desertification in Southwest China: Impacts, causes, and restoration. Earth-Science Reviews. 2014b, 132, 1–12. 10.1016/j.earscirev.2014.01.005

43. Rumpel, C., Alexis, M., Chabbi, A., Chaplot, V., Rasse, D.P., Valentin, C., &Mariotti, A. Black carbon contribution to soil organic matter composition in tropical sloping land under slash and burn agriculture. Geoderma. 2006, 1301-2, 35–46. 10.1016/j.geoderma.2005.01.007

44. Brodowski, S., Amelung, W., Haumaier, L., &Zech, W. Black carbon contribution to stable humus in German arable soils. Geoderma. 2007, 1391-2, 220–228. 10.1016/j.geoderma.2007.02.004

45. Llorente, M., Glaser, B., &Turrión, M.B. Storage of organic carbon and black carbon in density fractions of calcareous soils under different land uses. Geoderma. 2010, 1591-2, 31–38. 10.1016/j.geoderma.2010.06.011

46. Knicker, H. Pyrogenic organic matter in soil: Its origin and occurrence, its chemistry and survival in soil environments. Quaternary International. 2011, 2432, 251–263. 10.1016/j.quaint.2011.02.037

47. Glaser, B., Balashov, E., Haumaier, L., Guggenberger, G., &Zech, W. Black carbon in density fractions of anthropogenic soils of the Brazilian Amazon region. Organic Geochemistry. 2000, 317–8, 669–678. 10.1016/S0146-6380(00)00044-9

48. Ussiri, D.A., &Johnson, C.E. Characterization of organic matter in a northern hardwood forest soil by ^13^C NMR spectroscopy and chemical methods. Geoderma. 2003, 1111-2, 123–149. 10.1016/S0016-7061(02)00257-4

49. Li, G.X., Sun, L., Wang, J.Y., Dou, X., Ji, S.Z., Hu, T.X., &Gao, C.Y. Effects of pyrogenic carbon addition after fire on soil carbon mineralization in the Great Khingan Mountains peatlands Northeast China. Science of The Total Environment. 2023, 864, 161102. 10.1016/j.scitotenv.2022.161102

50. Yang, W.Q., Deng, R.J., &Zhang J. Forest litter decomposition and its responses to global climate change. Chinese Journal of Applied Ecology. 2007, 1812, 2889–2895.https://www.cjae.net/CN/Y2007/V18/I12/2889

51. Sun, J.B., Gao, F., Song, J.F., &Chui, X.Y. Distributions of soil particulate organic carbon and black carbon of two forest types in changbai mountain. Forest Research. 2017, 302,222–231. 10.13275/j.cnki.lykxyj.2017.02.006

52. Zhang, L.Q., &Zhang, M.K. Effects of land use on particulate organic carbon and black carbon accumulation in red and yellow soils. Chinese Journal of Soil Science. 2006, 374, 662–665. 10.3321/j.issn:0564-3945.2006.04.009

53. Dai, T., Li, A.F., &Zhang, M.K. Distribution characteristics of black carbon in agricultural soils of northern zhejiang plain. Chinese Journal of Soil Science. 2009, 406, 1321–1324. CNKI:SUN:TRTB.0.2009-06-023

54. Li, S.X., Yin, Y.F., Yang, Y.S., Gao, R., Ma, H.L., &Li, F.F. Effects of black carbon addition on soil labile organic carbon and native soil organic carbon. Soils. 2013, 451, 79–83. 10.3969/j.issn.0253-9829.2013.01.012

55. Edmondson, J., Stott, I., Potter, J., Lopez-Capel, E., Manning, D., Gaston, K., &Leake, J. Black carbon contribution to organic carbon stocks in urban soil. Environmental science & technology. 2015, 4914, 8339–8346. 10.1021/acs.est.5b00313

56. López-Martín, M., Velasco-Molina, M., &Knicker, H. Variability of the quality and quantity of organic matter in soil affected by multiple wildfires. Journal of Soils and sediments. 2016, 16, 360–370. 10.1007/s11368-015-1223-2

57. Wang, Q., Liu, M., Yu, Y.P., Du, F.F., &Wang, X. Black carbon in soils from different land use areas of Shanghai, China: level, sources and relationship with polycyclic aromatic hydrocarbons. Applied geochemistry. 2014b, 47, 36–43. 10.1016/j.apgeochem.2014.04.011

58. Liu, Z.Y., &Zhang, M.K. Components of organic carbon pool and profile distribution of δ^13^C values in typical stagnic anthrosols from hangzhou-jiaxing-huzhou plain. Journal of Zhejiang University Agriculture & Life Sciences. 2010, 363, 275–281. 10.3785/j.issn.1008-9209.2010.03.006

59. Das, O., Wang, Y., &Hsieh, Y.P. Chemical and carbon isotopic characteristics of ash and smoke derived from burning of C3 and C4 grasses. Organic Geochemistry. 2010, 413, 263–269. 10.1016/j.orggeochem.2009.11.001

60. Wang, P., Zhou, W., Niu, Z., Cheng, P., Wu, S., Xiong, X., &Du, H. Emission characteristics of atmospheric carbon dioxide in Xi’an, China based on the measurements of CO_2_ concentration, 14C and δ13C. Science of the Total Environment. 2018, 619, 1163–1169. 10.1016/j.scitotenv.2017.11.125

